# Recurrent SARS-CoV-2 mutations at Spike D796 evade antibodies from pre-Omicron convalescent and vaccinated subjects

**DOI:** 10.1101/2023.08.22.554362

**Authors:** Evan A Elko, Heather L Mead, Georgia A Nelson, John A Zaia, Jason T Ladner, John A Altin

**Author notes:** these authors contributed equally (Evan A Elko, Heather L Mead).

## Abstract

SARS-CoV-2 lineages of the Omicron variant rapidly became dominant in early 2022 and frequently cause human infections despite vaccination or prior infection with other variants. In addition to antibody-evading mutations in the Receptor Binding Domain, Omicron features amino acid mutations elsewhere in the Spike protein, however their effects generally remain ill-defined. The Spike D796Y substitution is present in all Omicron sub-variants and occurs at the same site as a mutation (D796H) selected during viral evolution in a chronically-infected patient. Here we map antibody reactivity to a linear epitope in the Spike protein overlapping position 796. We show that antibodies binding this region arise in pre-Omicron SARS-CoV-2 convalescent and vaccinated subjects, but that both D796Y and D796H abrogate their binding. These results suggest that D796Y contributes to the fitness of Omicron in hosts with pre-existing immunity to other variants of SARS-CoV-2 by evading antibodies targeting this site.

## Introduction

The Omicron variant of SARS-CoV-2 was first identified in Botswana and South Africa in November 2021 and rapidly spread across the globe to become the predominant circulating strain. Omicron is associated with a higher reinfection rate and reduced vaccine efficacy (1,2), and is distinguished by a striking number of mutations compared to previous variants (3,4) – which individually and collectively have the potential to alter the virus’s transmissibility, pathogenicity, and ability to escape immune responses. Of particular interest are a core set of persistent mutations shared by all circulating Omicron sub-variants: these are enriched in the Spike protein that mediates viral entry, and particularly in the Receptor Binding Domain (RBD) within its S1 subunit, whose interaction with host ACE2 is required for infection (5). Recent studies have established that many of these Omicron RBD mutations greatly diminish the binding and/or neutralizing effects of antibodies raised by prior variants (6–8), explaining their emergence and persistence. However, Omicron also features conserved mutations outside of the RBD, including in the S2 subunit that enables membrane fusion and entry into the host cell. The consequences of these mutations remain less well-understood.

The persistent Omicron Spike mutation D796Y is located in the N-terminal part of the S2 subunit immediately upstream of the fusion peptide. D796Y may confer structural advantages by enhancing the interaction between the Spike-TMPRSS2 complex (9) and potentially by altering the presentation of a nearby immunogenic glycan epitope (10). Intriguingly, D796Y occurs at the same residue as a mutation (D796H) that emerged independently in 2020 in a chronically infected patient treated with remdesivir and convalescent plasma, which was shown to confer reduced sensitivity to neutralization by convalescent plasma (11). These results suggest that D796Y may have a similarly evasive effect, which is supported with a recent preprint showing that S2 mutations, including D796Y, can reduce the neutralization potency of S1-binding antibodies (12).

In this study, we test the hypothesis that exposure to pre-Omicron Spike proteins induces D796-binding antibodies that are evaded by both the D796H and D796Y mutations. We first use public sequence data to chart the emergence of D796 mutations throughout the pandemic, and find evidence consistent with a role in conferring increased viral fitness, independently of other Omicron mutations. We then use a multiplexed and sensitive peptide-based assay to identify a public antibody epitope overlapping position D796 in cohorts with pre-Omicron infection or vaccination, and to quantify the effects of the D796H and D796Y mutations on these antibody:epitope interactions.

## Results

### Phylogenetic analysis of mutations at D796 throughout the SARS-CoV-2 pandemic

The occurrence of D796Y in all lineages of the SARS-CoV-2 Omicron variant could reflect either a positive effect on viral fitness (e.g. through antibody evasion and/or an increase in transmissibility; adaptive hypothesis) or simply physical linkage with other fitness-increasing mutations (neutral hypothesis). To begin to distinguish these possibilities, we looked for evidence for independent occurrences of D796 mutations across the SARS-CoV-2 phylogeny. We downloaded >6 million publicly available SARS-CoV-2 sequences and identified clusters containing AA mutations at position 796 and >10 descendents (**Figure 1A**). The frequency of mutations identified in the phylogenetic tree were compared to the theoretical frequency of AA mutations resulting from all possible single nucleotide non-synonymous substitutions at codon 796. This analysis revealed that the D796Y and D796H substitutions have independently arisen at least 35 and 14 times throughout the pandemic, respectively, with the earliest documented instances preceding the emergence of Omicron by up to 13 months. Whereas the combined theoretical proportion of D>Y or D>H at this codon is 0.25, we observed these substitutions at a proportion >0.9 in pre-Omicron samples (p-value < 0.0001, based on 10,000 random simulations of single nucleotide non-synonymous substitution) (**Figure 1B**). These findings are consistent with previous analyses (13,14) and the hypothesis that D796Y confers a fitness advantage that is independent of other Omicron mutations.

**Figure 1.**
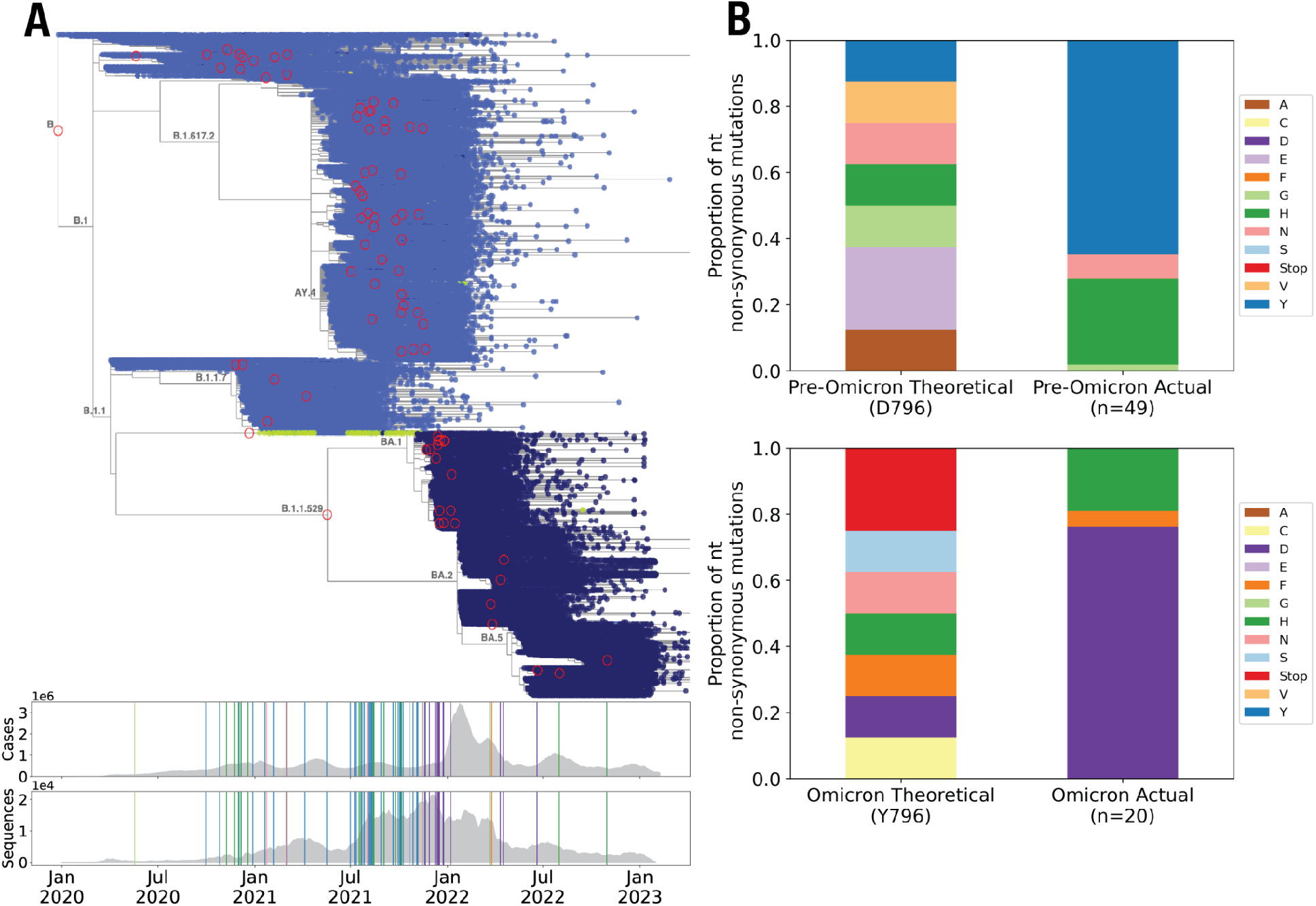
Phylogenetic analysis reveals that the D796Y and D796H substitutions occurred recurrently prior to the Omicron wave. **A**. Upper: Time-resolved phylogenetic tree including 6,329,365 SARS-CoV-2 sequences (Taxonium.org). The color of the points indicate the amino acid residue at position 796 of the SARS-CoV-2 Spike: Light Blue=D, Green = H, Dark Blue = Y. Red circles highlight nodes with mutations at 796 that were found in ≥10 descendants (a threshold set to minimize the impact of technical artifacts). Branch labels indicate PANGO lineage (27). Lower: Plots showing daily worldwide SARS-CoV-2 cases and total sequences per day present in the phylogenetic tree over a 2 year timescale (x-axis aligned with the phylogenetic tree in the upper panel). Vertical lines indicate the positions of mutants highlighted with red circles in the phylogenetic tree and colors show the mutation type (matching the bar graphs in (B)). **B**. The proportion of theoretically possible (left) compared to observed (right) non-synonymous amino acid substitutions resulting from single nucleotide mutations in the codon for D796 in pre-Omicron sequences (upper) and Y796 in post-Omicron sequences (lower).

### Infection and vaccination induce antibodies that bind a linear epitope overlapping D796

Inspection of the 3-dimensional structure of the Spike trimer indicates that D796 resides on a surface-exposed loop that is theoretically available for antibody binding (**Supplemental Figure 1**). To test whether pre-Omicron antibody responses recognize epitopes overlapping position D796, we mapped linear epitopes at high resolution across the complete ancestral SARS-CoV-2 Spike protein by applying a highly-multiplexed binding assay (‘PepSeq’) to plasma from participants who were vaccinated against or convalescent from SARS-CoV-2 infection prior to the emergence of Omicron. The PepSeq platform has been previously described (15–17) and enables highly-multiplexed serological analysis using programmable libraries of DNA-barcoded peptide probes that are synthesized from DNA templates in massively-parallel *in vitro* reactions. The SARS-CoV-2 (SCV2) PepSeq library (16) comprises 2,500 tiled 30mer peptides of which 1,244 and 390 represent the ancestral Spike and Nucleocapsid proteins, respectively, with each peptide overlapping its neighbor by 29 amino acids. We used SCV2 to assay plasma from two cohorts sampled in 2020-2021 prior to the emergence of the Omicron variant: (i) a cohort of unvaccinated donors who were either previously infected (53 subjects) or naive (58 subjects) (“convalescent cohort”), and (ii) a cohort of 21 SARS-CoV-2 naive subjects sampled prior to and after vaccination with mRNA-1273 (“vaccinated cohort”).

To identify epitopes recognized in response to infection or vaccination, we compared reactivity between infected and naive subjects in the convalescent cohort and matched pre- and post-vaccination samples in the vaccinated cohort (**Figure 2A**). This analysis revealed robust antibody responses to infection and vaccination: we detected significantly increased reactivity in 172/79 and 209/0 Spike/Nucleocapsid peptides in the convalescent and vaccinated cohorts, respectively (hereafter termed “responding peptides”). The restriction of Nucleocapsid responding peptides to the convalescent cohort is expected, consistent with the lack of this protein sequence in the mRNA-1273 vaccine. Importantly, we observed strong responses to peptides overlapping position D796 in both cohorts: among the 30 assayed peptides tiled across this position, 19 and 25 were significantly elevated in the convalescent and vaccinated cohorts, respectively (**Figure 2B,C**), with up to 2.3 fold-increases in enrichment Z scores. These response frequencies among peptides overlapping D796 (0.63 and 0.83 in convalescent and vaccinated subjects, respectively) are substantially higher than the corresponding rates among the overall set of 1,244 Spike peptides (0.14 and 0.17).

**Figure 2.**
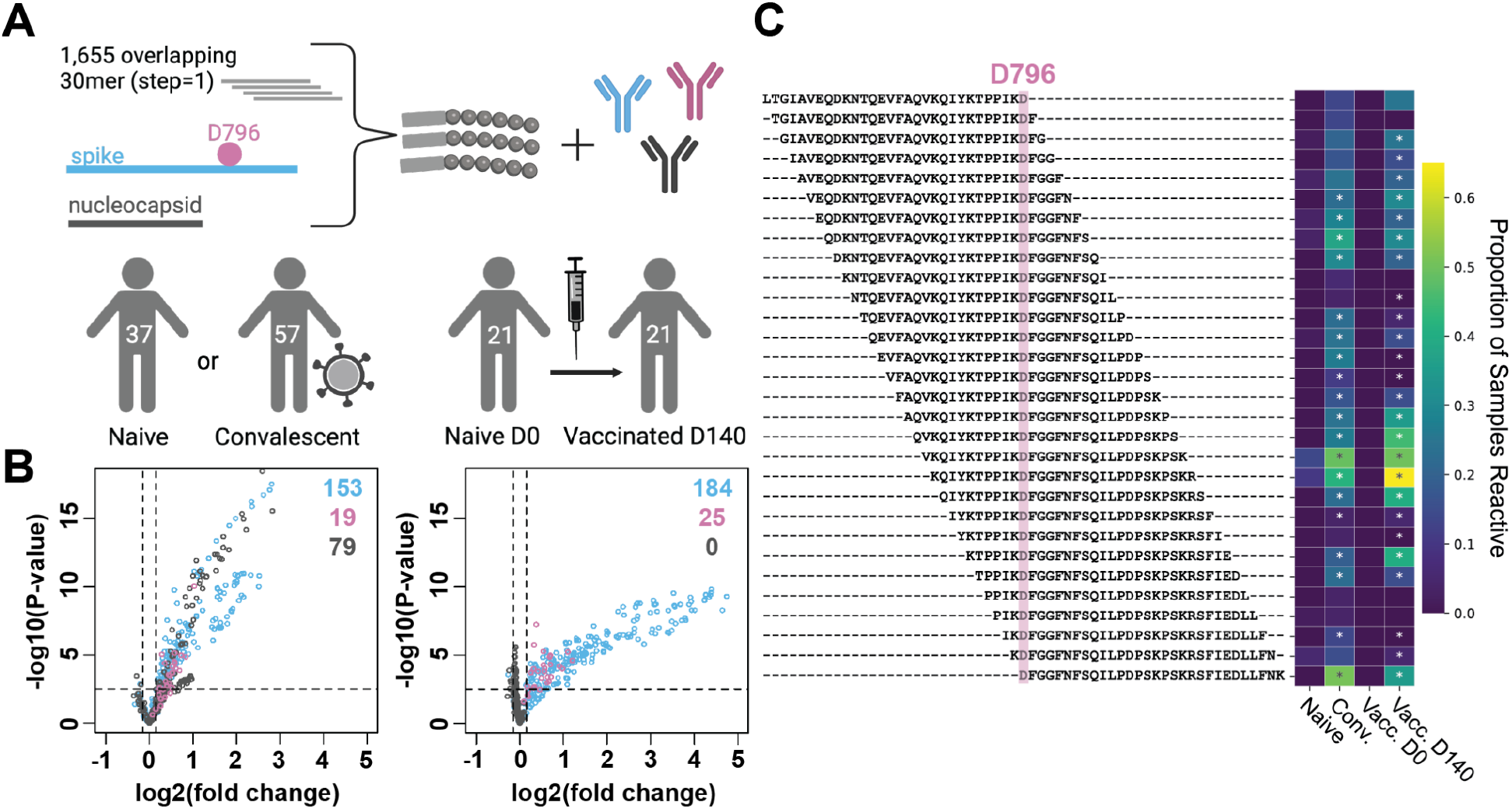
Antibody responses to D796-containing Spike peptides following natural infection with or vaccination against SARS-CoV-2. **A**. A library containing 1,655 DNA-barcoded peptides tiling the complete ancestral SARS-CoV-2 Spike (S) and Nucleocapsid (N) proteins at high resolution (30mers overlapping by 29 amino acids) was synthesized using the PepSeq platform. This library was used to measure IgG reactivity at epitope resolution in naive, convalescent and vaccinated subjects (n=115 total participants). **B**. Volcano plots showing the magnitude and significance of responses targeting other S peptides (blue), S peptides overlapping D796 (pink), and N peptides (gray), in comparisons of naive versus convalescent participants (left) and naive versus vaccinated timepoints (right). Dashed lines indicate thresholds at log2(fold changes) of ±0.15 and -log10(p-value) of 2.5. **C**. Heatmap showing the proportion of participants who were reactive to each peptide (at a Z-score threshold ≥ 6) in the indicated cohort across all 30 peptides that contained D796 (y-axis). Asterisks indicate peptides whose responses exceed the cohort-wide thresholds shown in B.

The positions of responding peptides within the Spike protein were highly similar between the convalescent and vaccinated cohorts (**Supplemental Figure 2**) and closely match previously-described epitope regions, including the fusion peptide and stem helix regions against which broadly-neutralizing antibodies have been described (18–20).

### Quantifying the impact of D796 mutations on antibody binding

Having identified antibody reactivity in pre-Omicron immune subjects to a region of the Spike protein that overlaps D796, we next sought to test the hypothesis that the D796Y and D796H mutants evade these responses. We quantified antibody binding to 3 versions of this position – WT, D796Y and D796H – in the pre-Omicron immune cohorts described above. To accomplish this, we generated a new PepSeq library in which we represented each of the 3 versions of position 796 in the form of 10 overlapping individual 30mer peptides (**Figure 3A**).Binding analysis of these mutated peptides in subjects selected according to their reactivity to the region containing position 796 (18 convalescent subjects at days 27-314 after testing positive for SARS-CoV-2 and 17 vaccinated subjects at day ∼140 post-vaccination) revealed a marked and consistent reduction in antibody reactivity to peptides containing either the D796Y or D796H substitutions in both cohorts, compared to the unmutated versions (**Figure 3B,C**). These effects were broadly consistent across each of the 10 tiled peptides for each mutation (**Supplemental Figure 3**). We conclude that mutation of D to either Y or H evades antibody reactivity to a linear epitope overlapping position 796 induced by exposure to ancestral Spike proteins.

**Figure 3.**
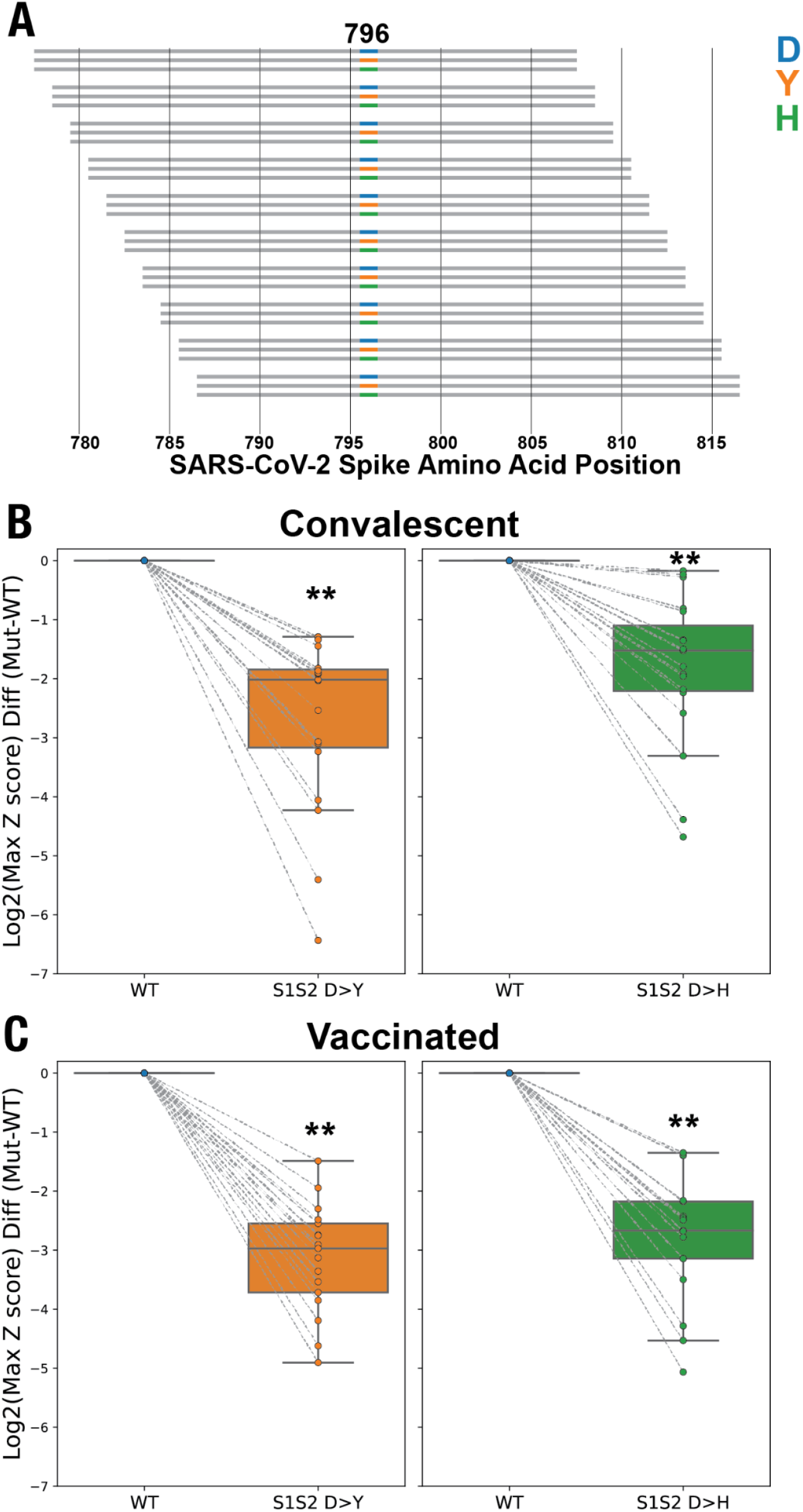
D796Y and D796H mutations abrogate the binding of infection- and vaccine-induced antibodies to Spike peptides. **A**. A PepSeq library was generated to compare 3 different residues at Spike position 796 – D (WT), Y and H – each replicated in 10 tiled overlapping peptides (total 30 peptides containing position 796). **B**,**C**. Reactivity differences between mutant (Y796, H796) and WT (D796) peptides in convalescent participants (upper, n=18) and vaccinated participants at day ∼140 (lower, n=17). Y-axes show maximum reactivity among the 10 overlapping peptides containing the indicated amino acid at position 796, normalized in each case to the WT signal from the same participant. ** = p-value < 0.0001 by Wilcoxon Signed-Rank Test.

## Discussion

In this study, we map antibody binding to an epitope that overlaps residue D796 of SARS-CoV-2 Spike: a residue that has mutated to Y and H recurrently throughout the pandemic, including in a well-characterized chronically-infected patient (D796H), and is now fixed in all Omicron subvariants (D796Y). By studying convalescent and vaccinated cohorts, we show that this epitope is frequently targeted by the polyclonal antibody response to ancestral Spike proteins, but that antibody binding is abrogated by both the D796H and D796Y mutations. Together, these results reveal a novel immunological consequence of these important Spike mutations, and raise the possibility that immune pressure mediated by antibodies binding to this epitope in the Spike S2 subunit has helped to shape the evolution of SARS-CoV-2.

Although we observe antibodies binding the D796-containing epitope in a high fraction of subjects (>70% of convalescent/vaccinated subjects – **Figure 2C, Supplemental Figure 2B**), their functional consequences remain to be determined. Previous work indicated that D796H reduces sensitivity to neutralization by convalescent plasma, however this mutation alone also causes an opposing and marked intrinsic infectivity defect that is unrelated to antibodies (11). In light of our observation that D796Y and D796H both evade antibodies to the linear epitope containing position 796 to similar degrees (**Figure 3B,C**), we hypothesize that D796Y may likewise confer a partial neutralization-evading effect. Mechanistically, the proximity of this epitope to the S2’ cleavage site raises the possibility that such antibodies could neutralize SARS-CoV-2 by inhibiting S2’ cleavage by TMPRSS2, an event that is necessary for host cell infection (21). Alternatively, or additionally, these antibodies may act against SARS-CoV-2 through neutralization-independent mechanisms.

In addition to the class of antibodies defined here, D796 mutations may evade antibodies binding Spike at distal sites through complex conformational changes, as indicated by the recent observation that D796Y decreases sensitivity to neutralizing antibodies binding the S1 subunit (12). This reveals an intriguing potential for multi-epitope evasion by a single mutation, although the relative contributions of these effects to overall immune evasion remain to be determined.

By identifying a population of antibodies whose binding depends on D796, our results suggest immune pressure as an important driver of the emergence of D796 mutations, which would be consistent with its role in shaping the evolution of SARS-CoV-2 more broadly. A more complete understanding of the mutational mechanisms by which SARS-CoV-2 has evaded, and continues to evade, immune responses may enable better prediction of its future evolution, as well as the identification of evasion-resistant vulnerabilities that can be exploited in the search for next generation therapies and vaccines.

## Methods

### Phylogenetic analysis

The phylogenetic tree of 6,329,365 publicly available SARS-CoV-2 sequences from GenBank (i.e. the INSDC databases), the China National Center for Bioinformation, and from COG-UK, along with estimated tip dates were downloaded from https://taxonium.org/?backend=https://api.cov2tree.org on Feb. 8, 2023. If no date was supplied for a given sequenced genome, tip dates were estimated using Chronumental(22). These tip dates were then used to calculate the number of sequences per day in the dataset. The number of global SARS-CoV-2 cases per day were downloaded from ourworldindata.org/covid-cases on Feb. 9, 2023. Taxonium(23) was used to visualize the phylogenetic tree and identify branches with mutations at D796 that had greater than 10 descendents. Nodes meeting this criteria were then used to calculate the proportion of single nucleotide non-synonymous amino acid mutations pre- and post-omicron becoming the dominant global variant. Theoretical distribution of single nucleotide non-synonymous amino acid mutations were calculated by considering all possible amino acid outcomes of single nucleotide mutations at the D796 codon. A simulation approach was utilized to determine the likelihood that the observed AA mutation frequency was caused by random chance based on the resulting theoretical AA distribution of single nucleotide non-synonymous substitutions. Ten-thousand iterations were run using the theoretical distributions pre and post-Omicron. For each iteration the number of observed mutations pre and post-Omicron (n=49 and n=20 respectively) were chosen from the theoretical distribution and the proportion of D>Y/H mutations was calculated and compared to the actual observed proportion of D>Y/H mutations.

### Study subjects

Under an IRB-approved study (IRB#20204, NCT04497779), unvaccinated COVID-19 convalescent individuals were recruited from participating clinical sites. Participants all experienced mild COVID-19 symptoms and donated blood within 7-142 days of their initial PCR-based diagnosis. Age, sex, positive or negative to SARS-CoV-2 spike IgG antibody, and sample collection dates are listed in Supplemental Table 1. Pre-pandemic naive control serum samples (n=24) were collected by Creative Testing Solutions (Phoenix, AZ) during January 2015 from multiple locations in California (Supplemental Table 1). The use of all pre-pandemic samples was reviewed by the TGen and NAU Research Compliance offices and determined not to be human subjects research.

The vaccinated cohort studied here has been previously described (15,16). Under an IRB-approved study (WIRB#1299650), 21 healthy participants were recruited from a local research institution and the surrounding community. Subjects donated blood prior to their first dose (baseline) of the Moderna COVID-19 vaccine (mRNA-1273) and then approximately 140 days from baseline (“day 140”). The characteristics and exact collection timepoints for each donor are listed in Supplemental Table 2.

### PepSeq libraries

The SCV2 PepSeq library contained 2,500 unique 30mer peptides and was designed to provide high-resolution coverage of the SARS-CoV-2 Spike and Nucleocapsid proteins, as previously described (16). Briefly, this library included all unique 30 amino acid peptides contained in consensus versions of the SARS-CoV-2 Spike and Nucleocapsid proteins generated early in the COVID-19 pandemic. This included 1244 Spike and 390 Nucleocapsid peptides, each of which was represented by three different nucleotide encodings.

The 796 mutant library is a derivative of our human virome version 2 PepSeq library, which has been described in detail previously (15)). It contains 15,000 unique 30mer peptides including a set of 30 peptides designed to test the impact of the mutations at position 796 of the SARS-CoV-2 Spike: each of three residues (D, Y, H) was represented in 10 overlapping positions (tiled with a step of 1 amino acid, as shown in **Supplemental Figure 3A**).

### PepSeq library synthesis and antibody-binding assay

The SCV2 and 796 mutant library designs were encoded as libraries of 7,500 or 15,000 DNA oligonucleotides, respectively. These oligonucleotides were used to synthesize a corresponding ‘PepSeq’ library of DNA-barcoded peptides for multiplexed analysis of antibody reactivity profiles, as previously described (24). Briefly, the oligonucleotide library was PCR-amplified and then used to generate mRNA in an *in vitro* transcription reaction. The product was ligated to a hairpin oligonucleotide adaptor bearing a puromycin molecule tethered by a PEG spacer and used as a template in an *in vitro* translation reaction. Finally, a reverse transcription reaction, primed by the adaptor hairpin, was used to generate cDNA, and the original mRNA was removed using RNAse. To perform serological assays, the resulting DNA-barcoded peptide library was added to diluted plasma and incubated overnight. The binding reaction was applied to pre-washed protein G-bearing beads, washed, eluted, and indexed using barcoded DNA oligos. Following PCR cleanup, products were pooled, quantified, and sequenced using an Illumina NextSeq instrument resulting in the generation of between 47,383 to 3,917,807 reads per sample.

### PepSeq data analysis

PepSeq sequencing data was processed and analyzed as previously described (16) using PepSIRF v1.6.0 (25,26), as well as custom scripts (https://github.com/LadnerLab/PepSIRF/tree/master/extensions). First, the reads were demultiplexed and assigned to peptides using the PepSIRF *demux* module, allowing for one mismatch in each index sequence and two mismatches in the variable DNA tag region. The PepSIRF *norm* module was then used to normalize counts to reads per million (RPM). RPM normalized reads from seven buffer-only negative control assays were used to create bins for Z-score calculation using the PepSIRF *bin* module. To normalize for different starting peptide abundances within each bin, reads were further normalized by subtracting the average RPM from the buffer only controls (--diff option in *norm* module). Z-scores were calculated using the PepSIRF *zscore* module using the 70%(SCV2) or 60% (796 mutant library) highest density interval within each bin. The peptide Z-scores were compared between naive/convalescent and naive/vaccinated using a student’s t-test and peptides were considered significantly responsive if they had a log2(fold change) ≥ 0.15 and a -log10(p value) ≥ 2.5. The proportion of samples in each cohort that contained an enriched peptide (Z-scores ≥6) was calculated for all peptides overlapping the SARS-CoV-2 spike protein position 796.

## Supplemental Figures

**Supplemental Figure 1.**
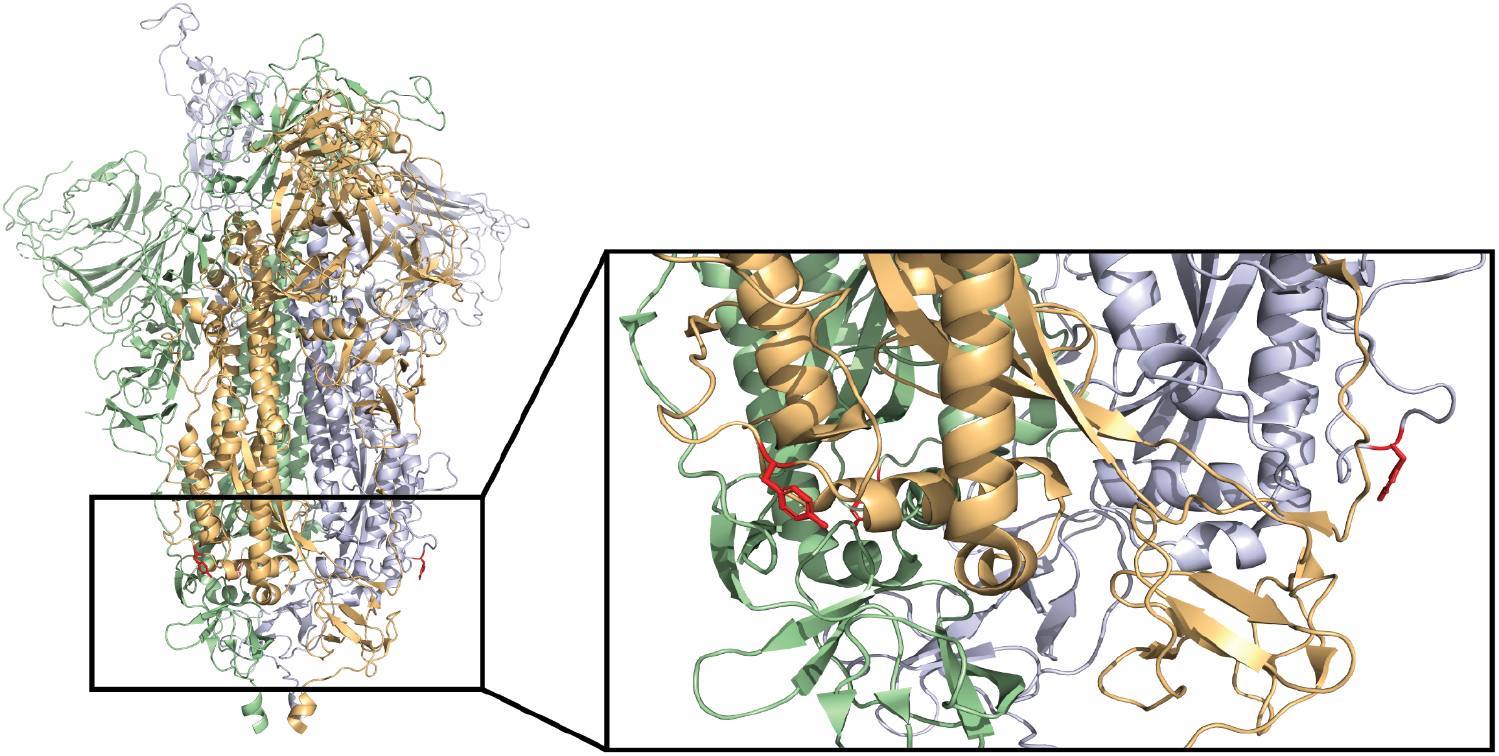
3D structure of Omicron Spike protein in the pre-fusion conformation highlighting Y796. Pre-fusion 3D model of Omicron SARS-CoV-2 Spike protein trimer (PDB:7TGW) with Y796 highlighted in red. Pastel orange, green and blue colors indicate the three Spike protomers.

**Supplemental Figure 2.**
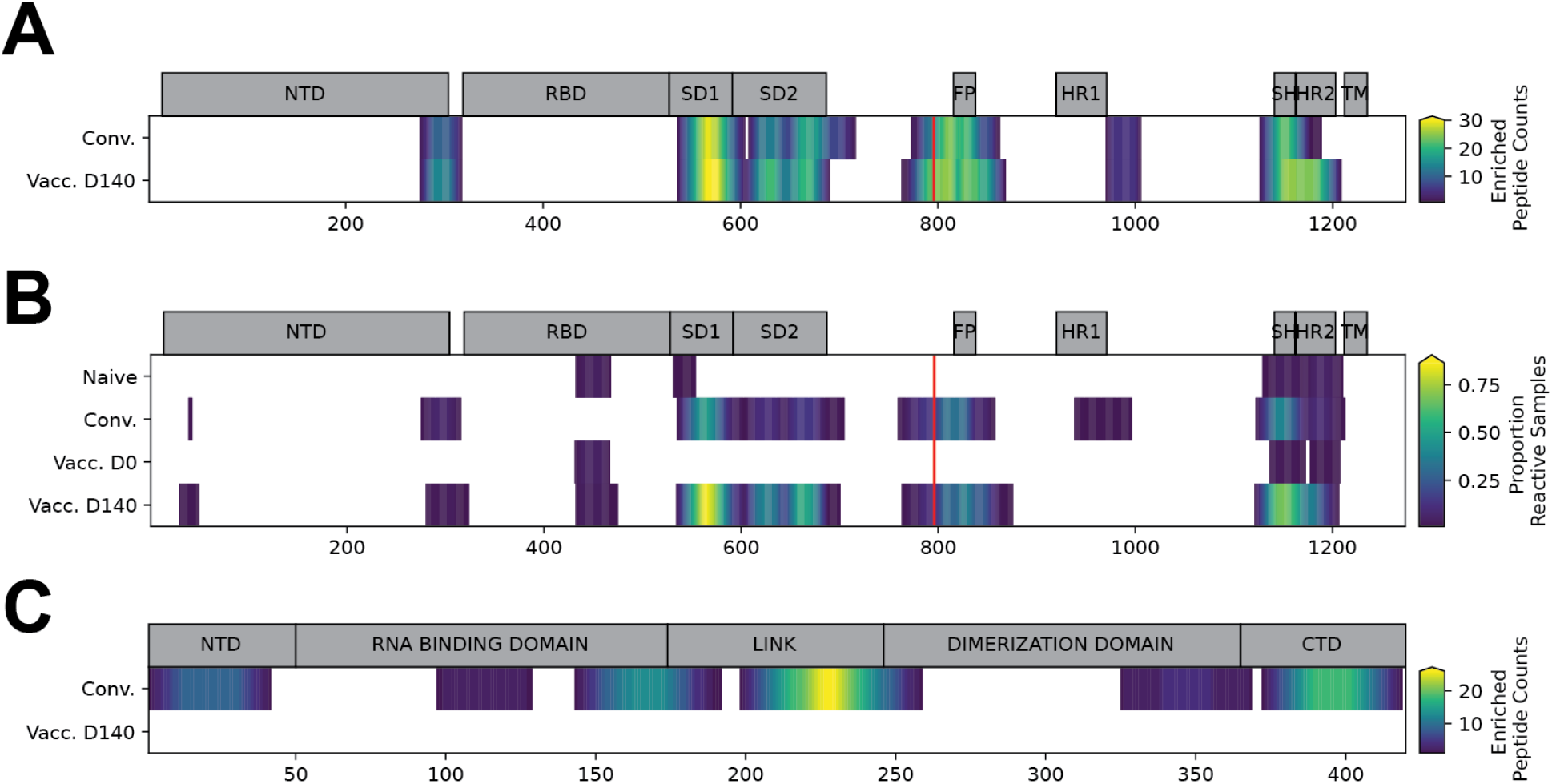
Spike and Nucleocapsid protein-wide linear epitope reactivity in convalescent and vaccinated subjects detected using PepSeq. Heatmaps enumerating enriched peptides overlapping each AA position across the entire SARS-CoV-2 Spike (**A**) and Nucleocapsid (**C**) proteins. Enriched peptides were determined by PepSeq Z-score comparisons between Naive and Convalescent (*upper*) or Naive and Vaccinated (day 140, *lower*) as in Figure 2b. **B**. Heatmap showing the proportion of samples with reactivity (Z-score ≥ 10) at each position within the SARS-CoV-2 spike protein, calculated by taking the mean proportion of reactive samples across all peptides that cover each amino acid position. The red lines in panels A and B indicate position 796.

**Supplemental Figure 3.**
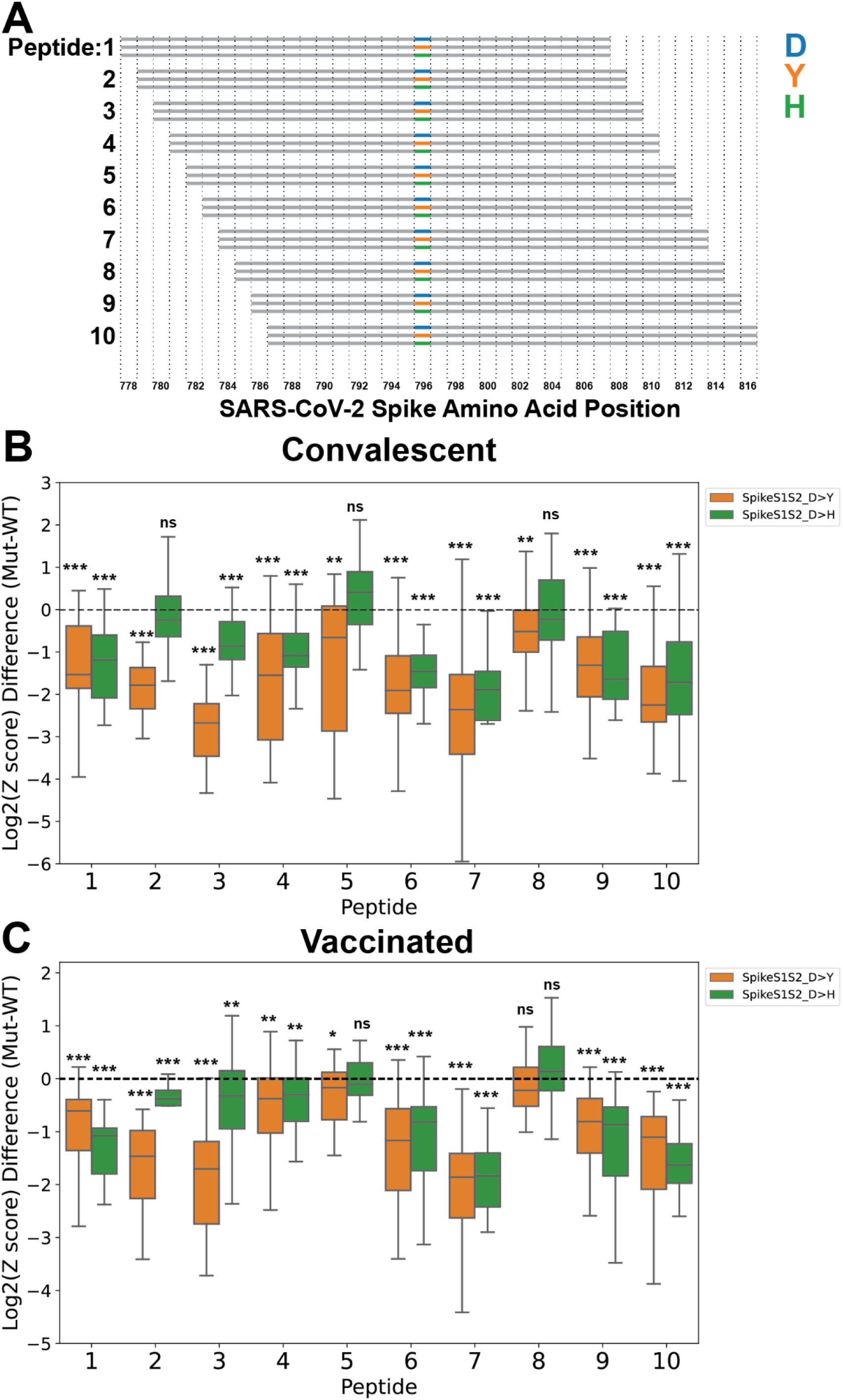
Reactivity patterns for individual wild-type and mutant peptides tiled across Spike position 796. **A**. Graphical representation of peptide tiling strategy around position 796. For each step there are three peptides, indicated by blue, orange or green that are identical except for position 796 where they contain a D (Ancestral SARS-CoV-2: blue), Y (Omicron: orange), or H (recurrent mutation, observed in chronic infection: green). X-axis indicates AA position within the SARS-CoV-2 spike protein and numbering on the Y-axis corresponds to the peptide identifiers used on the X-axis in B and C. Reactivity in **B** convalescent (n=18) and **C** vaccinated (n=17) subjects selected for reactivity at position 796, showing signal for each mutant peptide (Y796, H796), normalized to WT (D796) across the 10 tiled peptides described in A. * = p-value < 0.05, ** = p-value < 0.01, *** = p-value < 0.001 by individual t-test (Convalescent), or paired t-test (Vaccinated).

## References

1. Pulliam JRC, van Schalkwyk C, Govender N, von Gottberg A, Cohen C, Groome MJ, Dushoff J, Mlisana K, Moultrie H. Increased risk of SARS-CoV-2 reinfection associated with emergence of Omicron in South Africa. Science (2022) 376:eabn4947.

2. Cele S, Jackson L, Khoury DS, Khan K, Moyo-Gwete T, Tegally H, San JE, Cromer D, Scheepers C, Amoako DG, et al. Omicron extensively but incompletely escapes Pfizer BNT162b2 neutralization. Nature (2022) 602:654–656.

3. Tegally H, Wilkinson E, Giovanetti M, Iranzadeh A, Fonseca V, Giandhari J, Doolabh D, Pillay S, San EJ, Msomi N, et al. Detection of a SARS-CoV-2 variant of concern in South Africa. Nature (2021) 592:438–443.

4. Bansal K, Kumar S. Mutational cascade of SARS-CoV-2 leading to evolution and emergence of omicron variant. Virus Res (2022) 315:198765.

5. Huang Y, Yang C, Xu X-F, Xu W, Liu S-W. Structural and functional properties of SARS-CoV-2 spike protein: potential antivirus drug development for COVID-19. Acta Pharmacol Sin (2020) 41:1141–1149.

6. Cao Y, Wang J, Jian F, Xiao T, Song W, Yisimayi A, Huang W, Li Q, Wang P, An R, et al. Omicron escapes the majority of existing SARS-CoV-2 neutralizing antibodies. Nature (2022) 602:657–663.

7. Wang Q, Guo Y, Iketani S, Nair MS, Li Z, Mohri H, Wang M, Yu J, Bowen AD, Chang JY, et al. Antibody evasion by SARS-CoV-2 Omicron subvariants BA.2.12.1, BA.4 and BA.5. Nature (2022) 608:603–608.

8. Arora P, Kempf A, Nehlmeier I, Schulz SR, Jäck H-M, Pöhlmann S, Hoffmann M. Omicron sublineage BQ.1.1 resistance to monoclonal antibodies. Lancet Infect Dis (2023) 23:22–23.

9. Cheng Y, Zheng D, Zhang D, Guo D, Wang Y, Liu W, Liang L, Hu J, Luo T. Molecular recognition of SARS-CoV-2 spike protein with three essential partners: exploring possible immune escape mechanisms of viral mutants. J Mol Model (2023) 29:109.

10. Gobeil SM-C, Henderson R, Stalls V, Janowska K, Huang X, May A, Speakman M, Beaudoin E, Manne K, Li D, et al. Structural diversity of the SARS-CoV-2 Omicron spike. Mol Cell (2022) 82:2050–2068.e6.

11. Kemp SA, Collier DA, Datir RP, Ferreira IATM, Gayed S, Jahun A, Hosmillo M, Rees-Spear C, Mlcochova P, Lumb IU, et al. SARS-CoV-2 evolution during treatment of chronic infection. Nature (2021) 592:277–282.

12. Kumar S, Delipan R, Chakraborty D, Kanjo K, Singh R, Singh N, Siddiqui S, Tyagi A, Jha S, Thakur KG, et al. Mutations in S2 subunit of SARS-CoV-2 Omicron spike strongly influence its conformation, fusogenicity and neutralization sensitivity. bioRxiv (2023)2023.03.05.531143. doi: 10.1101/2023.03.05.531143

13. Maher MC, Bartha I, Weaver S, di Iulio J, Ferri E, Soriaga L, Lempp FA, Hie BL, Bryson B, Berger B, et al. Predicting the mutational drivers of future SARS-CoV-2 variants of concern. Sci Transl Med (2022) 14:eabk3445.

14. Zhao LP, Lybrand TP, Gilbert PB, Payne TH, Pyo C-W, Geraghty DE, Jerome KR. Rapidly identifying new coronavirus mutations of potential concern in the Omicron variant using an unsupervised learning strategy. Sci Rep (2022) 12:19089.

15. Elko EA, Nelson GA, Mead HL, Kelley EJ, Carvalho ST, Sarbo NG, Harms CE, Le Verche V, Cardoso AA, Ely JL, et al. COVID-19 vaccination elicits an evolving, cross-reactive antibody response to epitopes conserved with endemic coronavirus spike proteins. Cell Rep (2022)111022.

16. Ladner JT, Henson SN, Boyle AS, Engelbrektson AL, Fink ZW, Rahee F, D’ambrozio J, Schaecher KE, Stone M, Dong W, et al. Epitope-resolved profiling of the SARS-CoV-2 antibody response identifies cross-reactivity with endemic human coronaviruses. Cell Rep Med (2021) 2:100189.

17. Kelley EJ, Henson SN, Rahee F, Boyle AS, Engelbrektson AL, Nelson GA, Mead HL, Anderson NL, Razavi M, Yip R, et al. Virome-wide detection of natural infection events and the associated antibody dynamics using longitudinal highly-multiplexed serology. Nat Commun (2023) 14:1783.

18. Pinto D, Sauer MM, Czudnochowski N, Low JS, Tortorici MA, Housley MP, Noack J, Walls AC, Bowen JE, Guarino B, et al. Broad betacoronavirus neutralization by a stem helix-specific human antibody. Science (2021) 373:1109–1116.

19. Zhou P, Song G, Liu H, Yuan M, He W-T, Beutler N, Zhu X, Tse LV, Martinez DR, Schäfer A, et al. Broadly neutralizing anti-S2 antibodies protect against all three human betacoronaviruses that cause deadly disease. Immunity (2023) 56:669–686.e7.

20. Dacon C, Tucker C, Peng L, Lee C-CD, Lin T-H, Yuan M, Cong Y, Wang L, Purser L, Williams JK, et al. Broadly neutralizing antibodies target the coronavirus fusion peptide. Science (2022) 377:728–735.

21. Hoffmann M, Kleine-Weber H, Schroeder S, Krüger N, Herrler T, Erichsen S, Schiergens TS, Herrler G, Wu N-H, Nitsche A, et al. SARS-CoV-2 Cell Entry Depends on ACE2 and TMPRSS2 and Is Blocked by a Clinically Proven Protease Inhibitor. Cell (2020) 181:271–280.e8.

22. Sanderson T. Chronumental: time tree estimation from very large phylogenies. bioRxiv (2021) doi: 10.1101/2021.10.27.465994

23. Sanderson T. Taxonium, a web-based tool for exploring large phylogenetic trees. Elife (2022) 11:p doi: 10.7554/eLife.82392

24. Henson SN, Elko EA, Swiderski PM, Liang Y, Engelbrektson AL, Piña A, Boyle AS, Fink Z, Facista SJ, Martinez V, et al. PepSeq: a fully in vitro platform for highly multiplexed serology using customizable DNA-barcoded peptide libraries. Nat Protoc (2022) doi: 10.1038/s41596-022-00766-8

25. Fink ZW, Martinez V, Altin J, Ladner JT. PepSIRF: a flexible and comprehensive tool for the analysis of data from highly-multiplexed DNA-barcoded peptide assays. arXiv (2020) doi: arXiv:2007.05050

26. Brown AM, Bolyen E, Raspet I, Altin JA, Ladner JT. PepSIRF + QIIME 2: software tools for automated, reproducible analysis of highly-multiplexed serology data. (2022) doi: 10.48550/ARXIV.2207.11509

27. Rambaut A, Holmes EC, O’Toole Á, Hill V, McCrone JT, Ruis C, du Plessis L, Pybus OG. A dynamic nomenclature proposal for SARS-CoV-2 lineages to assist genomic epidemiology. Nat Microbiol (2020) 5:1403–1407.

